# The intrinsic cortical geometry of reading

**DOI:** 10.1101/2025.03.10.641861

**Authors:** R. Austin Benn, Alexander Holmes, Robert Scholz, Wei Wei, Francesco Alberti, Victoria Shevchenko, Ulysse Klatzmann, Rocco Chiou, Carla Pallavicini, Robert Leech, Jonathan Smallwood, Elizabeth Jefferies, Pierre-Louis Bazin, Daniel S. Margulies

## Abstract

How does the brain support the complex processes that allow us to read? Using predictive modeling we establish that visual and association cortex are closer together in individuals with stronger oral reading ability. These findings indicate that large-scale cortical geometry provides a scaffold that supports the coordinated processing required to read.

## Main

One of the unique capacities of human cognition is the ability to transform written words into meaningful representations of events and concepts through reading^1–3^. Reading is a core aspect of modern society^4–6^, and difficulties in this process lead to interpersonal, educational, and cultural challenges^7^. Reading engages two processing routes: a lexical route that facilitates rapid recognition of familiar words via the visual word form area, and a sublexical route that supports unfamiliar text via phonological association regions such as the superior temporal and supramarginal gyrus. Both routes of reading depend on the coordination of activity between distributed regions that span from visual and association cortex^2,4–7^.

Recently, it has been argued that the physical distance between regions of sensory and association cortex allows these regions to separate their functional roles^8^, enabling the brain to balance concrete and abstract descriptions of the outside world^9^. Our study set out to test this hypothesis by examining if the distance between cortical regions influences the efficiency with which we read. We measured pairwise cortical distance between parcels from the Schaefer 400 atlas^10^ using a discovery cohort comprising data from 996 participants from the Human Connectome Project Young Adult cohort^11^. We applied connectome predictive modeling^12^ to compare how word recognition and pronunciation relates to the physical structure of the cerebral cortex, measured as the distance between regions along the cortical surface. This cortical distance measure was compared with another geometric feature, parcel surface area, and with traditional resting-state functional connectivity based on time-series correlations between pairs of regions. Our index of reading ability was oral word reading — the capacity to recognize and pronounce words — as assessed with the Oral Reading Recognition Test, which uses irregularly- and regularly-spelled words to engage both lexical and sublexical reading routes^13^.

Our analysis began by selecting the features of brain architecture associated with oral reading ability (Fig. S1). For surface area, positive and negative associations with better oral reading reflect parcel expansion and contraction, respectively. For cortical distance, positive associations indicate longer inter-regional separation, and negative associations indicate shorter distances. For functional connectivity, positive and negative associations reflect stronger and weaker time-series correlations between regions. For each feature type, the positively and negatively associated features were summed and used to predict individual differences in reading performance via linear regression^12^. To compare how cortical distance, surface area, and functional connectivity each contribute to word recognition and pronunciation, we built 13 models and evaluated their performance through 10 repetitions of 10-fold cross-validation in our discovery cohort. We report results from the Schaefer 400 atlas, which outperformed other atlases for both cortical distance and functional connectivity (Figs. S2, S3). Of these models, nine were unimodal: six compared surface area, cortical distance, and functional connectivity using only their positive or negative features, and three joint models combined the positive and negative features of each measure. The remaining four models evaluated multimodal combinations of the joint feature sets for surface area, cortical distance, and functional connectivity. These multimodal models aimed to investigate how the brain’s physical and functional properties jointly shape our ability to recognize and pronounce words.

All models except those using negative surface area significantly predicted word recognition and pronunciation (Fig. 1a, Table S1). To compare models, we used paired permutation tests on the mean difference of cross-validation performance, controlling the false discovery rate^14^ (Fig. 1a,b, Table S2). For unimodal models, this analysis found the positive surface area models outperformed the positive functional connectivity models (r = 0.132 vs. 0.100; p = 0.016), the negative cortical distance models outperformed the negative surface area and negative functional connectivity models (r = 0.131 vs. 0.070; p = 0.008, and r = 0.131 vs. 0.085; p = 0.016), and the joint functional connectivity models outperformed the joint surface area models (r = 0.177 vs. 0.132; p = 0.023; Fig. 1a,b).

**Fig. 1.**
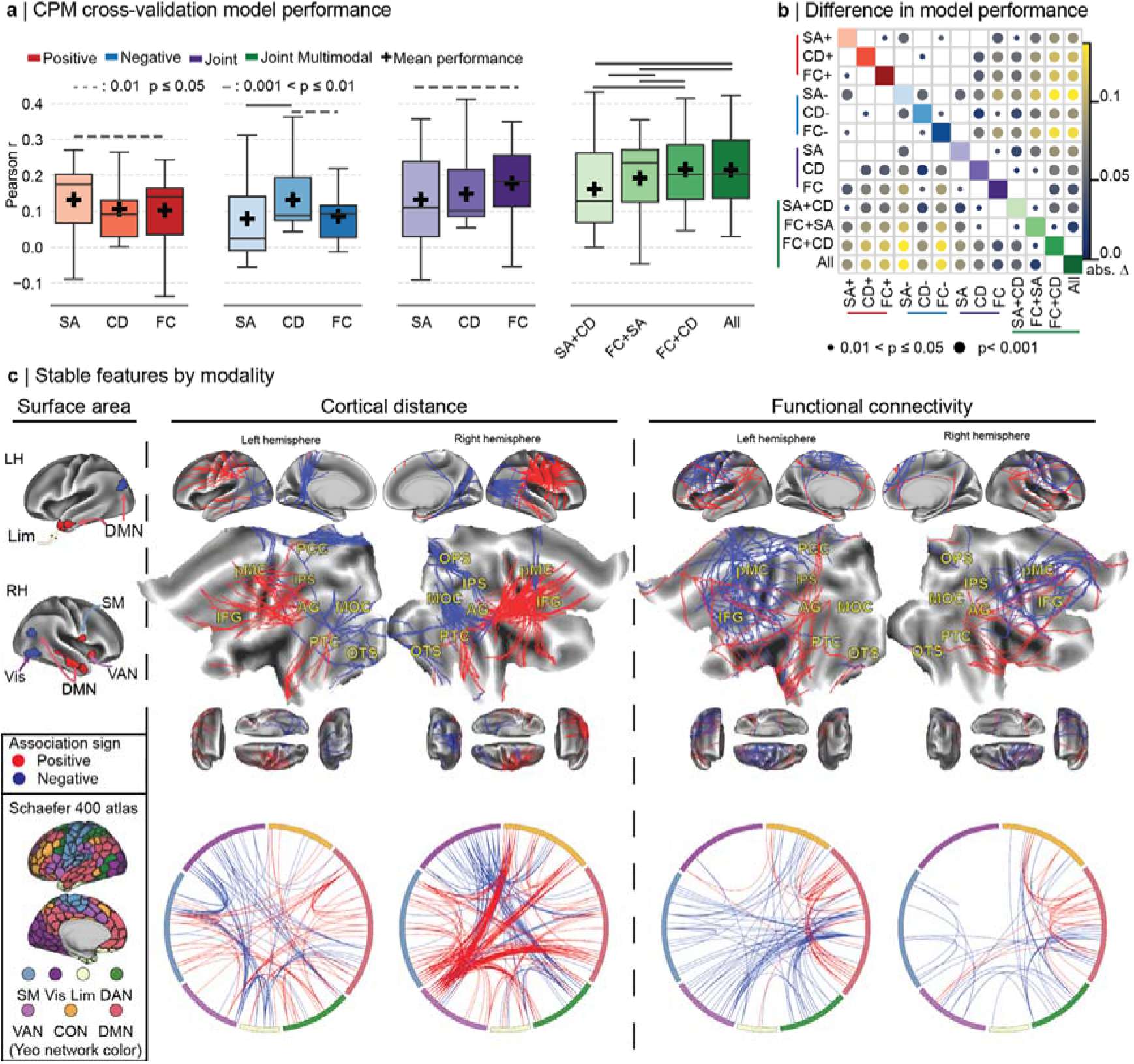
**a. Discovery cohort cross-validation performance of connectome predictive models:** Models using positively associated features (red) of surface area (SA), cortical distance (CD), and functional connectivity (FC) all predicted word recognition and pronunciation as measured by the Oral Reading Recognition Test. Negatively associated (blue) cortical distance and functional connectivity, and joint models combining positive and negative features of all three modalities (purple) predicted word recognition and pronunciation. Multimodal models (green) tested joint model combinations. For performance and p-values see Table S1. Boxes indicate the interquartile range with the median shown; whiskers extend to 1.5× IQR. **b. Pairwise difference in model performance:** Permutation testing identified significant differences in mean predictive performance between models across cross-validation folds. **c. Modality feature sets:** Stable feature sets for each modality, identified by consistent associations across all cross-validation folds. Highlighted areas in the cortical flat maps include middle occipital cortex (MOC), angular gyrus (AG), the posterior cingulate cortex (PCC), posterior temporal cortex (PTC), occipital temporal sulcus (OTS), occipitoparietal sulcus (OPS), intraparietal sulcus (IPS), premotor cortex (pMC), and inferior frontal gyrus (IFG). Chord plots of the CD and FC feature sets show pairwise relationships between parcels, organized by Yeo 7-network membership. All analyses used the Schaefer 400 parcellation^10^ . Network abbreviations include: Somatomotor (SM), Visual (Vis), Default Mode (DMN), Dorsal Attention (DAN), Ventral Attention (VAN), Control (CON), Limbic (Lim).

In the multimodal models, adding cortical distance to the joint surface area and joint functional connectivity models improved prediction performance (r = 0.132 vs. 0.161; p = 0.008; and r = 0.177 vs. 0.216; p = 0.008, respectively). Furthermore, adding surface area to the functional connectivity models improved prediction performance less than the addition of cortical distance (r = 0.192 vs. 0.216; p = 0.023). Adding surface area to the multimodal functional connectivity and cortical distance models produced no further improvement (r = 0.215 vs. 0.216; p = 1.0; Fig. 1a,b, Table S2). Together, these findings indicate that cortical distance provides complementary information that improves the prediction of word recognition and pronunciation over models built using only surface area and functional connectivity.

To characterize the brain features driving these predictions and enable external validation in an independent dataset, we built a stable feature set for each modality. These sets retained only those features that held consistent associations with word recognition and pronunciation across all cross-validation folds. In the surface area feature set, parcel surface areas positively associated with oral reading were found in the left temporal pole, right middle temporal gyrus, right insula, and right inferior central sulcus. Negatively associated parcel surface areas were located bilaterally in the lateral occipitoparietal boundary and in the right ventral-posterior temporal cortex (Fig. 1c).

The cortical distance feature set revealed two distinct spatial patterns: positively associated (longer) distances within association cortex, and negatively associated (shorter) distances between visual and association cortex (Fig. 1c)^1,15^. The positively associated distances primarily separated the ventral attention, executive, and default mode networks. Specifically, distances were greater between posterior temporal cortex and the temporal pole, and between lateral prefrontal cortex and the angular gyrus.

Negatively associated distances were distributed across both the lateral and medial surfaces of the cortex (Fig. 1c). They primarily positioned the visual cortex closer to posterior regions of the ventral and dorsal attention networks and the default mode network. This was observed on the lateral surface in both hemispheres for distances spanning the middle occipital and posterior temporal cortex, as well as between the posterior temporal cortex and angular gyrus. In the left hemisphere, we observed shorter distances between the angular gyrus and the middle occipital cortex, and in the right hemisphere the middle occipital cortex was closer to the superior temporal planum. Both these patterns were consistent with compression along the sensory-to-association axis.

Shorter distances in the medial cortex were lateralized, and in the left hemisphere the ventral visual cortex and precuneus were closer to the medial temporal pole. In the right medial cortex, one set of shorter distances positioned the medial temporal pole closer to the ventral-posterior visual cortex. A second set ran through the occipitoparietal sulcus, which positioned the lateral intraparietal sulcus closer to the medial temporal cortex.

Overall, the distance feature set is notable in that despite capturing two distinct ends of cortical organization, the positive and negative distances form part of known reading pathways^6^. Specifically, shorter distances bilaterally drew the angular gyrus and ventral occipitotemporal cortex toward visual cortex, with longer distances extending onward to the temporal pole (Fig. 1c).

Finally, the functional connectivity feature set aligned with prior research^5,16^, and was primarily distributed across association cortex with bilateral positive connectivity within and between the default mode and control networks, and negative connectivity between attention networks and the default mode network (Fig. 1c).

These feature sets were then used to test for generalization in an external validation cohort consisting of data from 396 participants from the Lifespan Human Connectome Project Aging cohort. We trained nine models using the stable positive, negative, and joint feature sets from each modality on the full discovery cohort. Models were then evaluated based on their out-of-sample prediction of word recognition and pronunciation in the validation cohort (Fig. 2a,b).

**Figure 2.**
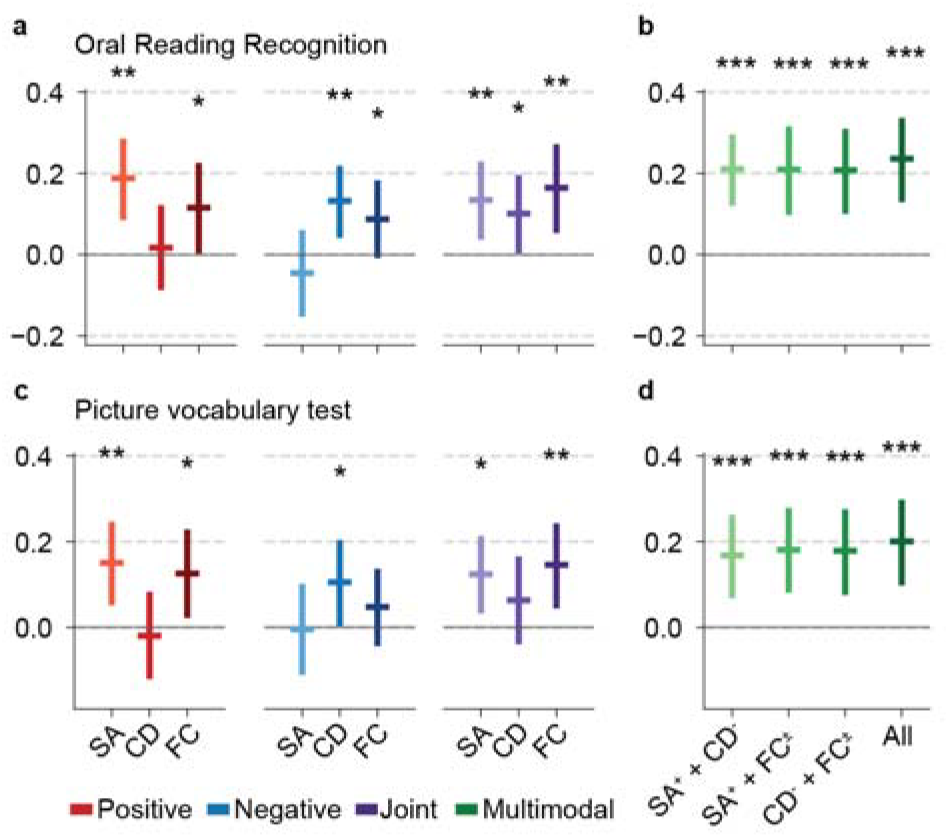
Model generalization to the validation cohort. Models were trained to predict word recognition and pronunciation as measured by the Oral Reading Recognition Test in the discovery cohort and evaluated for out-of-sample generalization in the external validation cohort. **a** Generalization of unimodal models to word recognition and pronunciation in the validation cohort. The positive surface area (SA) and functional connectivity (FC) models, the negative cortical distance (CD) model, and all joint unimodal models significantly predicted out-of-sample performance (Table S3). **b** Generalization of multimodal models to word recognition and pronunciation in the validation cohort. All multimodal models showed significant out-of-sample prediction (Table S3). **c** Out-of-sample and out-of-task generalization of unimodal models to word–picture association, measured using the Picture Vocabulary Test in the validation cohort. The positive and joint surface area and functional connectivity models, as well as the negative cortical distance model, significantly predicted performance (Table S4). **d** Out-of-sample and out-of-task generalization of multimodal models to word–picture association in validation cohort. All multimodal models showed significant predictive performance (Table S4). Markers show mean Pearson r; error bars denote 95% confidence intervals. Asterisks indicate significance of generalization from discovery to validation cohorts after correction for multiple comparisons using a two-step false discovery rate procedure, applied separately to unimodal and multimodal model families.

Across modalities, the strongest out-of-sample predictions came from positive surface area, negative cortical distance, and joint functional connectivity feature sets (Fig. 2a). We then used the best-performing feature sets to build multimodal models, excluding the negative surface area and positive cortical distance features which failed to generalize out of sample (Table S3). The resulting multimodal models reflected our cross-validation findings from the discovery cohort, as the addition of cortical distance consistently improved our ability to predict word recognition and pronunciation (Fig. 2b).

One limitation of the Oral Reading Recognition Test is that it assesses an individual’s ability to read words aloud but does not directly assess lexical-semantic access^13^. To assess if our models could do this, we tested their ability to predict word–picture association as measured by the Picture Vocabulary Test in our validation cohort^13,17^. Unlike the Oral Reading Recognition Test, this assessment was presented auditorily and did not require an oral response. As with reading aloud, the positive surface area, negative cortical distance, and joint functional connectivity models significantly predicted word–picture association (Fig. 2c, Table S4). Similarly, the addition of cortical distance to the surface area and functional connectivity models improved prediction (Fig. 2d). These findings suggest that the shorter distances identified in model training may partially capture the lexical access processes shared by both tasks, independent of pronunciation.

Although both shorter and longer distances predicted word recognition and pronunciation in the discovery cohort, only the shorter distances generalized to the validation cohort (Fig. 2a,c). Rather than being contradictory, this asymmetry suggests a single underlying effect: the longer distances converge spatially with the shorter ones, such that the impact of contraction between visual and association cortex extends to anterior regions along the lexical and sublexical reading pathways^6,18^. The longer distances may therefore be a geometric consequence of the same sensory-to-association compression, rather than an independent property, which is consistent with their failure to generalize. On this account, the stable, replicable signal is the proximity between visual and association cortex, which provides a physical scaffold that supports the transmission of visual information in orthographic processing^19^. This interpretation is further supported by the observation that the distributions of surface area and functional connectivity shift substantially across cohorts in parallel with oral reading ability, whereas the shorter distance feature set remains remarkably stable (Table S5), yielding nearly identical predictive effect sizes in cross-validation (r = 0.131) and external validation (r = 0.132). Instead, this geometric principle of sensory-association proximity in language appears to extend across sensory modalities. Consistent with this possibility, our models generalized to the auditorily presented Picture Vocabulary task (Fig. 2c,d), aligning our work with a recent study that found shorter distances between primary auditory cortex and auditory association areas are linked to enhanced acoustic–phonological processing^20^.

Broadly, our study highlights the importance of understanding how local areal expansion, regional brain activity, and the spatial arrangement of the cortex work together to shape cognition and behavior^21^. In this interplay, our findings reveal that the spatial arrangement of the cortex enhances our understanding of word recognition and pronunciation, complementing our ability to predict it from local areal expansion and functional brain activity (Figs. 1, 2). Initial investigations into how brain function and structure support complex behaviors have focused on understanding how local features relate to different features of cognition^22^. More recently, studies have established that the physical landscape of the cortex can explain brain activity during tasks^23^. In this context, our study bridges these views, extending the role of geometry as a stable physical scaffold that shapes not only brain activity^23^, but complex behaviors such as word recognition and pronunciation. Our study is thus a crucial next step in understanding how evolution has shaped both brain structure, and through it, the emerging landscape of brain function that supports the complex cognitive and behavioral repertoire that define us as a species.

## Methods

### Data

Our analyses used a discovery cohort comprising data from 996 participants in the Human Connectome Project Young Adult S1200 preprocessed data release^11^, and a validation cohort which used data from 396 participants from the Lifespan Human Connectome Project Aging cohort^24^. The discovery cohort was used in cross-validation (Fig. 1) and as the training set for external generalization (Fig. 2). The discovery cohort contained 463 males and 533 females, and had a mean age of 28.7 years (standard deviation 3.7). The validation cohort had 163 males and 233 females, with a mean age of 55.5 years (standard deviation 13.9). For both cohorts, participants were included if they had complete functional magnetic resonance imaging time-series data, and had no missing values for the Oral Reading Recognition Test, Picture Vocabulary Test, or the following confound variables: acquisition period, age, sex, diastolic and systolic blood pressure, education and income levels, intracranial volume, brain volume, and surface area of the left and right hemispheres.

A T1-weighted structural image was acquired with the following parameters: 0.7-mm isotropic resolution, TR = 2400 ms, flip angle of 8°, TE = 2.14 ms, FOV = 224 × 224 mm. The native surface was then constructed and resampled in the 32K_FS_LR space via Human Connectome Project minimal preprocessing pipelines^25^.

Functional connectivity measures were derived from functional magnetic resonance imaging data acquired on a 3 Tesla Siemens Connectome Skyra scanner. Four 15-minute resting state scans were acquired for each participant with the phase encoding alternated between right-to-left (RL) and left-to-right (LR) between each run. Acquisition parameters were as follows: 2.0-mm isotropic resolution, TR = 720 ms, multiband factor of 8, flip angle of 52°, TE = 33.1 ms, FOV = 208 × 180 mm, matrix size of 104 × 90, 72 slices, and echo spacing of 0.58 ms^25^.

### Parcellation schemes

Parcellation schemes define cortical areas and can differ substantially depending on the criteria used (e.g., functional connectivity or anatomical delineations)^10,26^. To assess the effect of parcellation choice on model performance and robustness, we first measured parcel surface area, cortical distance, and functional connectivity, and then built models from these features, evaluating their predictive performance with cross-validation in the discovery cohort (Figs. S2-S4). Parcellations tested included the Schaefer (2018) atlas at resolutions of 200, 400, 600, 800, and 1,000 parcels^10^; the Glasser atlas defined using combined anatomical and functional criteria with 360 regions^26^; and 400 randomly sampled equidistant points (200 per hemisphere).

### Parcel surface area

Parcel surface area was computed by first calculating vertex-wise surface areas on the Native 32k_fs_LR–aligned individual cortical surface using the Connectome Workbench command -surface-vertex-areas^27^. Vertex areas were then indexed by parcel, and the parcel surface area was recorded as the sum of vertex areas within a parcel for each parcellation (Fig. S4). In order to adapt this for the parcellation defined by random equidistant sampling, regions were grown using the zone_calc function in the surfdist package^28^, which assigns each vertex to the nearest sampled point as measured by geodesic distance.

### Cortical distance matrices

Cortical distance is measured along the cortical surface and remains constrained within each hemisphere. For this reason, the medial wall was excluded, and geodesic distance was measured independently for each hemisphere on the native midthickness surface resampled to the 32K_FS_LR using the surfdist package^28^. For each parcellation, we measured the shortest pairwise distance between the centroid of each parcel within a single hemisphere. This yielded an NxN pairwise distance matrix for each parcellation. From this matrix we extracted and vectorized the upper triangle of values, yielding a single vector of pairwise distances per hemisphere. The left and right hemisphere vectors were concatenated together for each participant which gave a single vector of pairwise distances per participant.

### Functional connectivity matrices

Time courses processed through the Human Connectome Project minimal preprocessing pipelines were used, and all four sessions were z-normalized and concatenated^25^. This yielded hour-long time series for each participant which were used to calculate the functional connectivity matrices. For each parcellation, we extracted the minimally preprocessed time series of each parcel by computing the average time course across its vertices. Pearson correlations were calculated for each pair of parcels within the same hemisphere which provided edgewise consistency to the cortical distance matrices. The resulting, within hemisphere connectivity matrices were then treated as in cortical distance; the upper triangle was extracted, vectorized, and the left and right hemisphere vectors were concatenated to yield a single vector of pairwise functional connectivity edges for each participant.

### Cross-validation in the discovery cohort

To estimate how reliably our models could predict performance on the Oral Reading Recognition Test, we split our dataset while respecting family structure in the discovery cohort with 10 repetitions of 10-fold cross-validation (CV). This resulted in 100 cross-validation folds, each containing approximately 891 training and 105 test participants. Cross-validation spanned both feature selection and predictive modeling (Fig. S1a).

### Data cleaning

The following variables were regressed out from the Oral Reading Recognition Test score and each entry in the surface area, cortical distance and functional connectivity feature vectors: *acquisition period, sex, age, diastolic and systolic blood pressure, education and income levels, intracranial volume, brain volume, and surface area of the left and right hemispheres*. Acquisition period was included as it accounts for reconstruction changes following Q1 of the Human Connectome Project. Sex was included as it also covaries with head size^29^ and was encoded as a binary variable. As we were not interested in the effects of age on reading, we also included it as a confound^30^. Age was preprocessed to enable downstream harmonization with the discovery cohort by computing its first two Legendre polynomials which yielded linear and orthogonal quadratic age terms. To avoid leakage, these terms were scaled using a fixed out-of-sample range of 18–100 years and applied consistently across both cohorts. Finally, blood pressure has also been associated with measures of functional connectivity^30^. For consistency, blood pressure was also regressed out for the surface area and cortical distance models. Measures estimating brain size (intracranial volume, brain volume, and the left and right cortical surfaces areas) were highly correlated with each other (Fig. S6). We therefore combined these variables into two brain-size components with principal component analysis (PCA), using the first two components that captured 98% of their original variance. PCA was performed exclusively on the training data; brain size–related features from test sets were projected into the learned PCA space during cross-validation and when testing generalization in the validation cohort. A final set of socioeconomic confounds (education and income) were also removed, given their correlation with reading ability (Fig. S6). Finally, prior to residualization, confounds and reading scores were transformed to approximate Gaussian distributions through a python port of the rank-based inverse normal transformation function in PALM^31^. Throughout data cleaning, transforms and residualizations were fitted exclusively on the training data and applied to the test data to avoid leakage. Cleaning was performed first on raw features during feature selection, and later the raw summary features prior to model fitting (Fig. S1b).

### Predictive modeling

Connectome-based predictive modeling consists of four stages: 1) Feature selection, 2) feature summarization, 3) model building, and 4) model evaluation (Fig. S1). We applied this approach to build models for each parcellation using parcel surface area, cortical distance and functional connectivity. For each parcellation and modality, we first correlated all available features to oral reading recognition scores in the training data, retaining those with an uncorrected p-value < 0.01. While subthreshold features may contribute meaningfully to prediction and, in some cases, improve prediction performance^32^, we retained this threshold to support a straightforward interpretation of cortical distance in its first application in connectome-based predictive modeling. Two models were constructed: (1) a model that retained the 50 strongest positive and 50 strongest negative correlations that stabilized the number of features selected across parcellations, and (2) a full model using all available features. The first model is presented in the main text; both are compared in Fig. S5.

For each set of selected features, we then split them by sign and summed their raw values, creating a positive and negative summary feature for each participant. These summary features were then cleaned as described above, removing the same set of confounds before they were mean and variance normalized. To avoid data leakage, confound removal and normalization were fit on the training set and learned transforms were applied to each test set in the cross-validation folds. These summary features were then used in linear models to predict performance on the Oral Reading Recognition Test (Fig. S1c).

For each modality (surface area, cortical distance, or functional connectivity), we trained three models per parcellation using the positive, the negative or both positive and negative (joint) summary features. In total, this yielded nine models per parcellation (Figs. S2-S5). Models were evaluated by calculating the Pearson correlation between the predicted and actual data in each fold’s test set. The overall performance was reported as the mean r-value obtained across all 100 folds. From these models, we observed that the Schaefer 400 parcellation using the top 50 features in each summary feature yielded the highest cross-validation performance; therefore, we selected this model for further analysis (Figs. S2-S5) using it to create four multimodal models using each modality’s joint model (Fig. 1a,b).

### Model significance

Model significance was assessed using a non-parametric permutation test. For each CV fold, we generated 1,000 permutations using a Human Connectome Project exchangeability block that takes into account family structure when shuffling this dataset^33^. Shuffling Oral Reading Recognition Test scores prior to feature selection broke the brain-behavior relationships, and the models were retrained and evaluated on their ability to predict the unshuffled test set. The p-value was calculated as the probability that the true model performance score was greater than that of the model trained on permuted data.

### Comparing performance across models

Our cross-validation scheme provided us with 100 estimates of model performance, i.e. one per cross-validation fold. To determine significant differences in model performance, we used a permutation test on the pairwise difference in mean performance between models. For each permutation, half of the fold-wise correlation values were randomly swapped between models 10,000 times, and the resulting distribution of mean performance differences was compared to the observed difference in mean correlations across all folds.

### External generalization to the validation cohort

#### Feature extraction and preprocessing

To test the generalization and robustness of our best-performing cortical distance and functional connectivity models (Schaefer 400; Figs. S2, S3), we evaluated them in a validation cohort using data from the Lifespan Human Connectome Project Aging cohort. We first established a stable set of features from our cross-validation study. Specifically, we selected surface area parcels, cortical distances, and functional connectivity edges whose sign remained consistent across all 100 cross-validation folds with a p-value below 0.01. For each modality and each individual, we summed the positive and negative feature values separately, yielding two summary features per modality.

Once the summary features were defined, confound removal was fitted on the full discovery cohort and then applied to the validation cohort, preventing data leakage and minimizing the influence of confound variables. We used the same procedure as in cross-validation to remove the effects of brain size, sex, blood pressure, education, and income. However, this was not possible for age, as the age ranges did not overlap between the discovery (24–34 years) and validation (35–90 years) cohorts. To account for this, after removing all other confounds and prior to model fitting and evaluation, age was regressed out within each cohort from the brain-derived predictive features only.

#### Significance testing of out-of-sample model transfer

Models were fit for each modality using the full discovery cohort and evaluated for non-chance predictive performance on the validation cohort with a permutation test. True behavior scores were shuffled 10,000 times to generate a null distribution of Pearson correlations between behavior and model predictions. This yielded a p-value that tests transfer of predictive signal from the discovery to the validation cohort; model performance comparisons were conducted exclusively within cross-validation in the discovery cohort. Reported p-values were corrected for multiple comparisons using a two-step Benjamini–Hochberg false discovery rate procedure^34^. Confidence intervals (95%) were estimated using bootstrap resampling (Fig. 2a,b, Table S3).

#### Construction of multimodal models

To assess whether combining modalities increased the out-of-sample point estimate of predictive performance, the features from the statistically significant model with the highest out-of-sample score from each modality were selected and combined to predict reading performance. Specifically, we used all combinations of the positive surface area, negative cortical distance, and joint functional connectivity features to build multimodal models. Generalization was assessed using permutation tests, consistent with the single-modality analyses. Confidence intervals were calculated using the same procedure. P-values for the multimodal models were corrected for multiple comparisons separately from the unimodal validation tests as they constitute a distinct, post-hoc test family conditional on the unimodal results.

#### Assessing feature and target drift

To assess changes in feature distributions and behavioral measures between the discovery and validation cohorts, we calculated Cohen’s *d* and the Wasserstein distance for the positive and negative halves of each feature set, as well as for oral reading ability (Table S5).

#### Out of task generalization

To test whether models trained to predict reading generalized to the Picture Vocabulary Test, models were trained to predict word recognition and pronunciation in the discovery cohort. The trained models were then frozen and applied to the validation dataset, where the target behavior was replaced with word–picture association. Significance of generalization for both unimodal and multimodal models was assessed using the same procedures described above (Fig. 2c,d, Table S4).

#### Ethics statement

All data analyzed in this study were obtained from the Human Connectome Project Young Adult and Lifespan HCP-Aging studies. Participant recruitment and informed consent procedures for the HCP Young Adult cohort were approved by the Washington University Institutional Review Board in accordance with the HCP protocols. The Lifespan HCP-Aging study obtained written informed consent from all participants; for those aged 80 years and older, capacity to provide informed consent was assessed using the MacArthur Capacity to Consent Scale, and only individuals who passed this assessment were enrolled. The present secondary analysis of these de-identified, openly shared data did not require further review.

## Data availability

The discovery cohort used in this study was sourced from the Human Connectome Project^25,26^. The validation cohort from the Lifespan Human Connectome Project Aging cohort^24^ accessed through the National Institute of Mental Health data archive (Data Access Request 23553). To facilitate replication, we have also provided the vectorized surface area, distance and functional connectivity matrices used in our analysis in a Zenodo repository at zenodo.org/records/20558695.

## Code availability

All code has been made publicly available in our GitHub repository at https://github.com/NeuroanatomyAndConnectivity/IntrinsicGeometryOfReading. This repository contains all scripts used for distance-based connectome predictive modeling, including feature selection, predictions, and the visualization of results.

## Acknowledgements

This research was funded by the European Research Council (ERC) under the European Union’s Horizon 2020 research and innovation programme (Grant agreement No. 866533) awarded to D.S.M. and supported by the NIHR Oxford Health Biomedical Research Centre (NIHR203316). The views expressed are those of the author(s) and not necessarily those of the NIHR or the Department of Health and Social Care. The Wellcome Centre for Integrative Neuroimaging is supported by core funding from the Wellcome Trust (203139/Z/16/Z and 203139/A/16/Z). JS was supported by (i) an award from the Government of Canada’s New Frontiers in Research Fund (NFRF) (grant ID: NFRF-2021-00183). All data were provided by the Human Connectome Project, WU-Minn Consortium (Principal Investigators: David Van Essen and Kamil Ugurbil; 1U54MH091657) funded by the 16 NIH Institutes and Centers that support the NIH Blueprint for Neuroscience Research; and by the McDonnell Center for Systems Neuroscience at Washington University.

## Author contributions

R.A.B. and D.S.M. conceptualized the study. R.A.B., P.-L.B., and D.S.M. developed the methodology. R.A.B. wrote the analysis code, performed the formal analysis, produced the visualizations, and wrote the original draft. A.H., R.S., W.W., F.A., V.S., U.K., R.C., C.P., R.L., J.S., and E.J. provided conceptual and methodological input. D.S.M. supervised the project and acquired funding. All authors reviewed and edited the manuscript and approved the final version.

