## Supplementary Information for "The intrinsic cortical geometry of reading"

**Supplementary Figures**

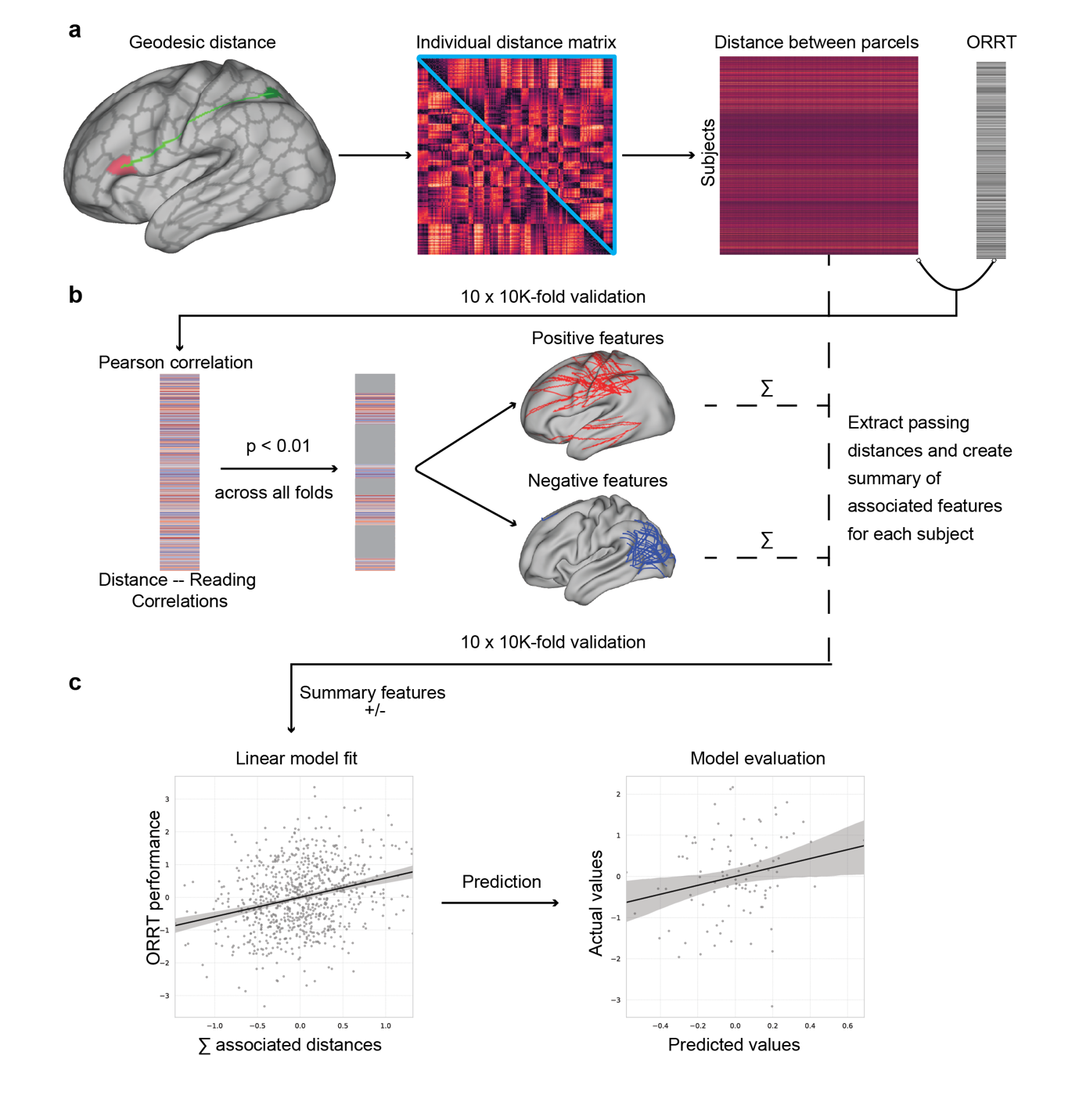

**Fig. S1. Adapting connectome-based predictive modeling to cortical distances**

**a** Geodesic distance was measured along the shortest path between all pairs of parcels from the Schaefer 400 atlas^10^. **b** Cortical distances associated with an uncorrected p-value < 0.01 were retained. The positive and negative associations were split, and summed separately for each individual in the dataset.^12^ **c.** The positive and negative features were used to fit a linear model and predict word recognition and pronunciation as measured by the Oral Reading Recognition Test (ORRT) using data from the discovery cohort.

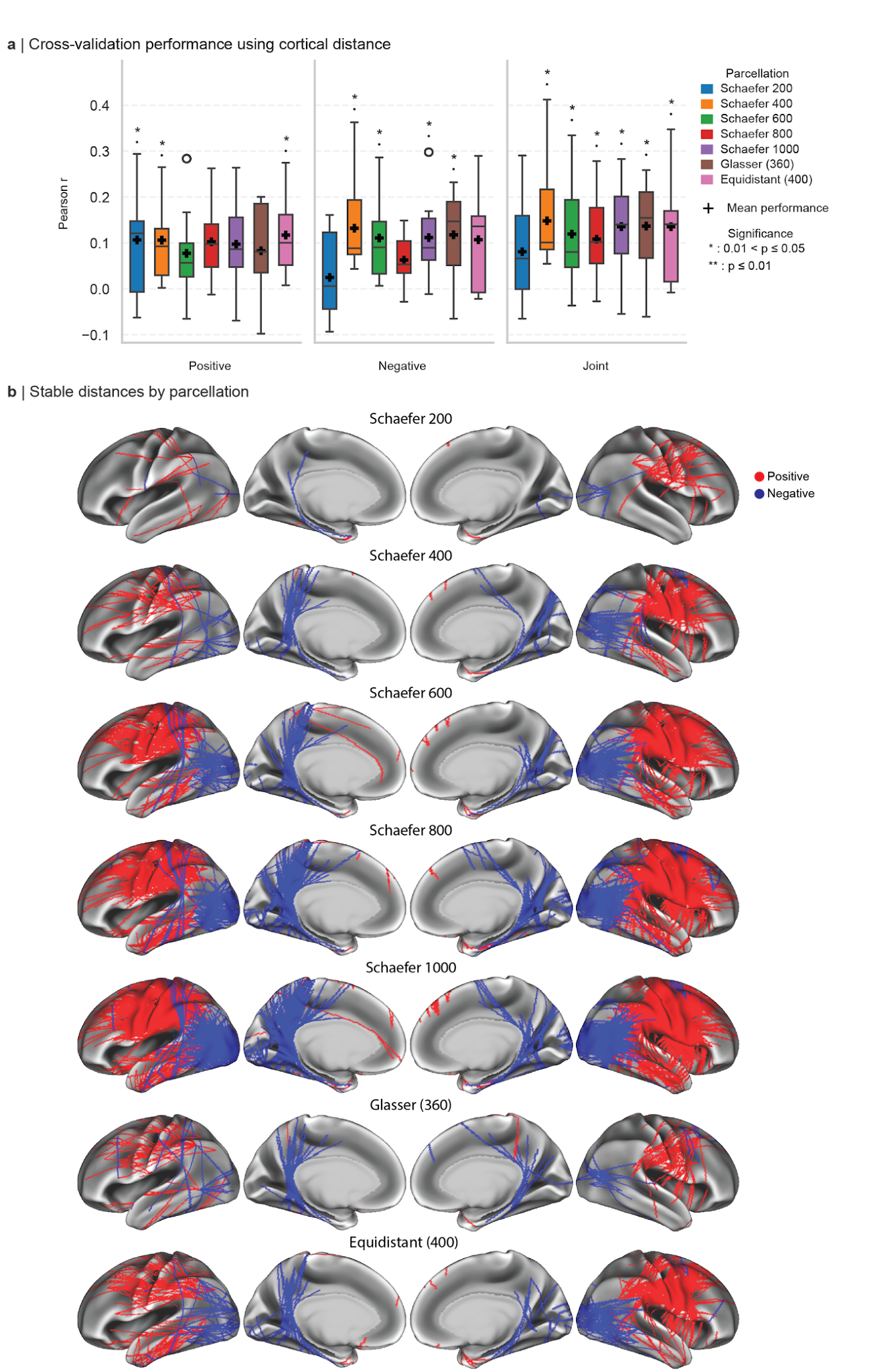

**Fig. S2 | Cortical distance model performance across parcellations. a** Cross-validation model performance using cortical distance measured from the Schaefer atlas at resolutions of 200, 400, 600, 800, and 1,000 parcels^10^; the Glasser atlas (360 regions)^26^; and 400 randomly sampled equidistant points (200 per hemisphere). Boxes indicate the interquartile range with the median shown; whiskers extend to 1.5× IQR. **b** Cortical distances with an uncorrected association p-value < 0.01 across all 100 cross-validation folds.

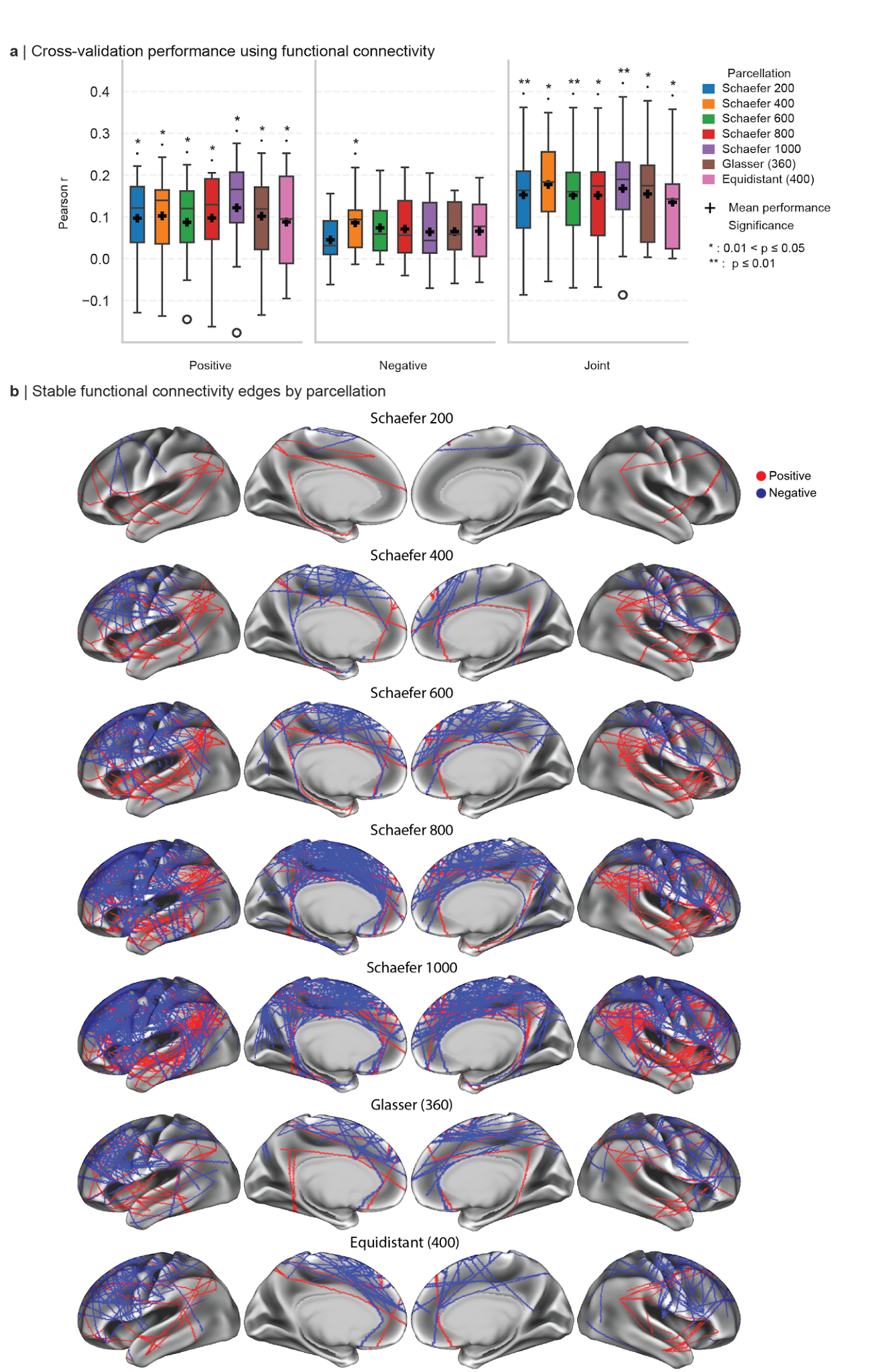

**Fig. S3 | Functional connectivity model performance across parcellations. a** Cross-validation model performance using within-hemisphere functional connectivity edges derived from the Schaefer atlas at resolutions of 200, 400, 600, 800, and 1,000 parcels^10^; the Glasser atlas (360 regions)^26^; and 400 randomly sampled equidistant points (200 per hemisphere). Boxes indicate the interquartile range with the median shown; whiskers extend to 1.5× IQR. **b** Functional connectivity edges with an uncorrected association p-value < 0.01 across all 100 cross-validation folds.

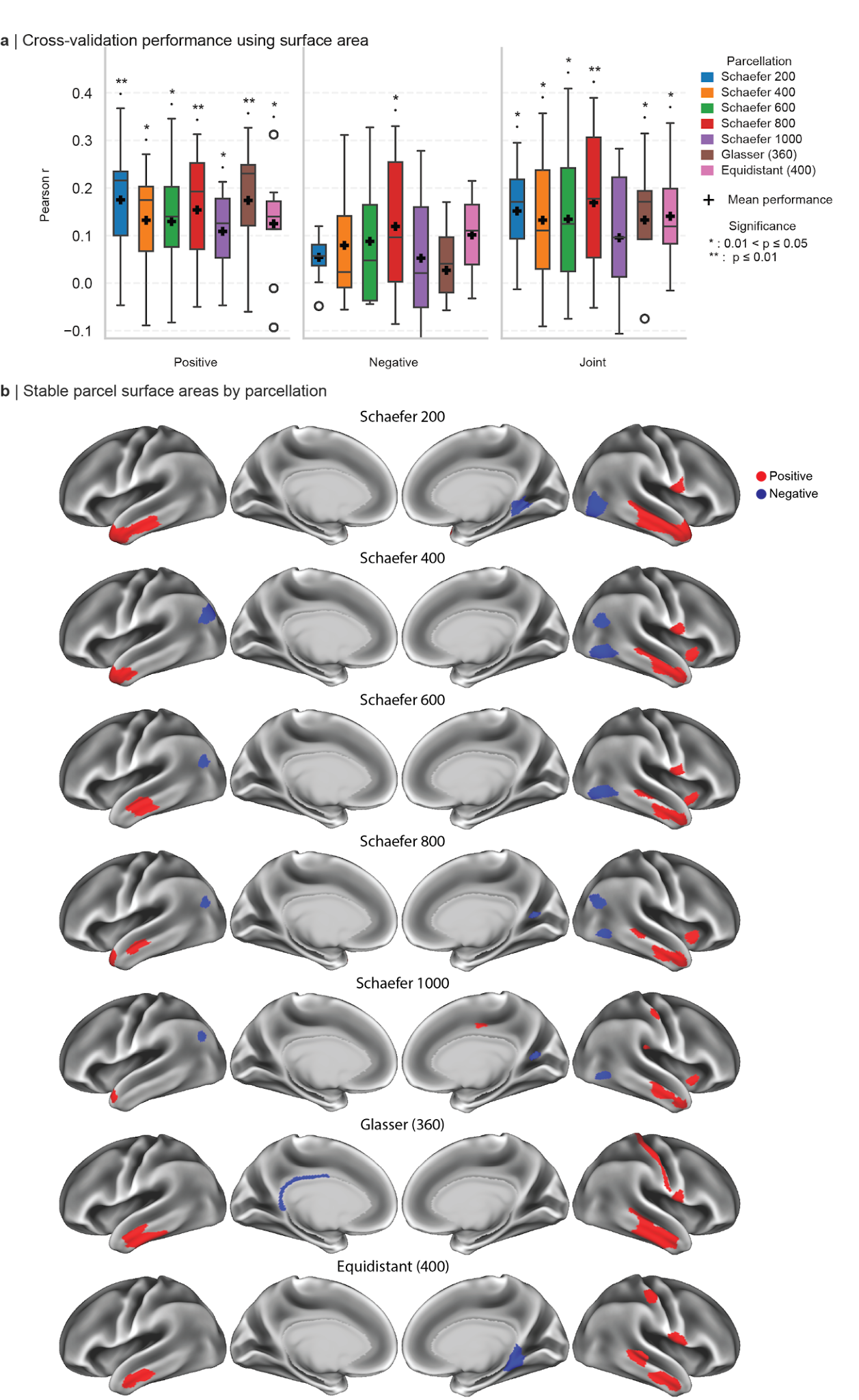

**Fig. S4 | Parcel surface area model performance across parcellations. a** Cross-validation model performance using parcel surface area derived from the Schaefer atlas at resolutions of 200, 400, 600, 800, and 1,000 parcels^10^; the Glasser atlas (360 regions)^26^; and 400 randomly sampled equidistant points (200 per hemisphere). Boxes indicate the interquartile range with the median shown; whiskers extend to 1.5× IQR. **b** Parcel surface areas with an uncorrected association p-value < 0.01 across all 100 cross-validation folds.

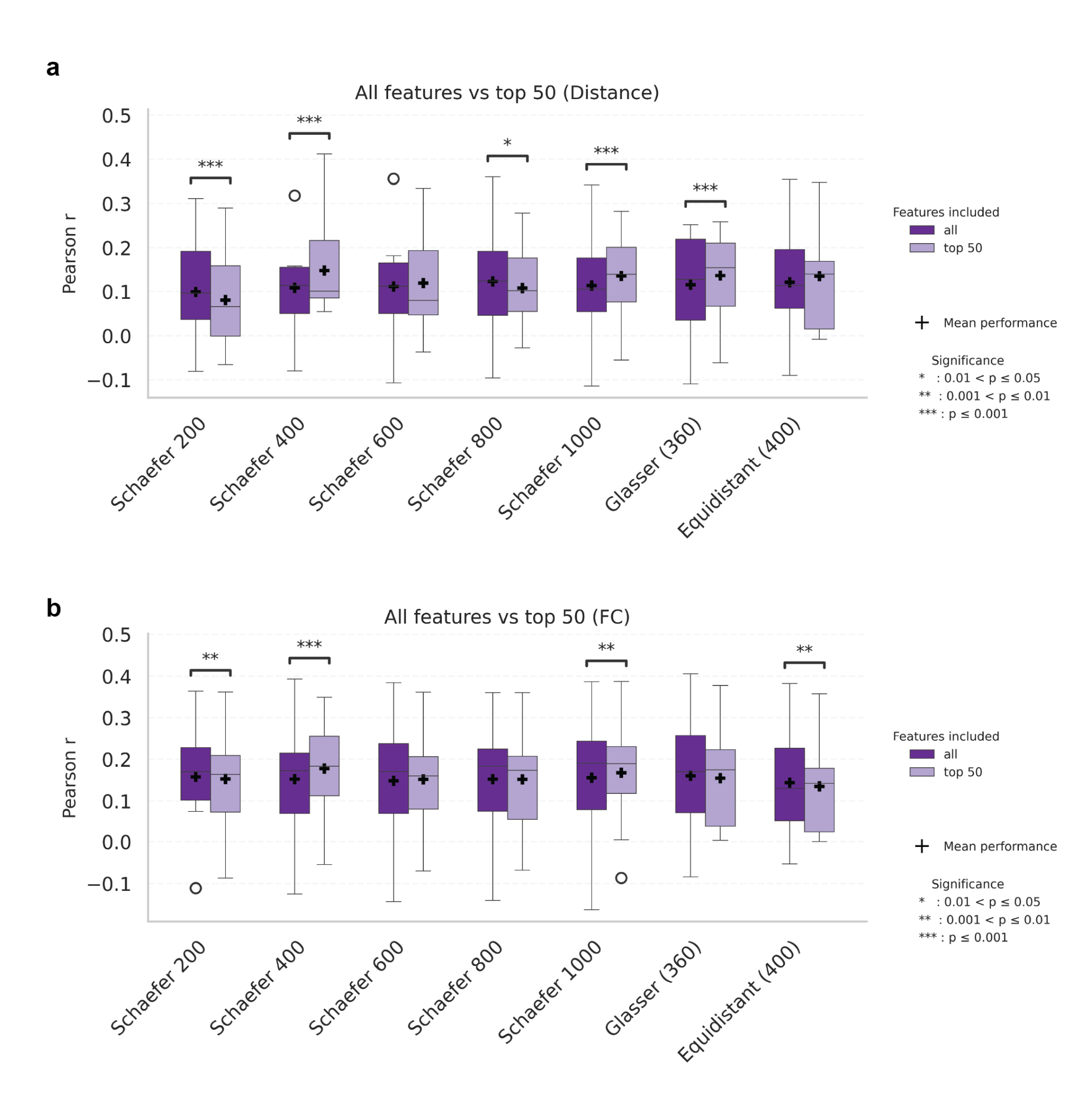

**Fig. S5 | Model performance: all features vs. top 50 most correlated features.** Cross-validation performance comparing models using all available features against models using only the top 50 most correlated features (50 positive, 50 negative) within each cross-validation fold. **a** Cortical distance (CD). **b** Functional connectivity (FC). Boxes indicate the interquartile range with the median shown; whiskers extend to 1.5× IQR. Crosses indicate mean performance.

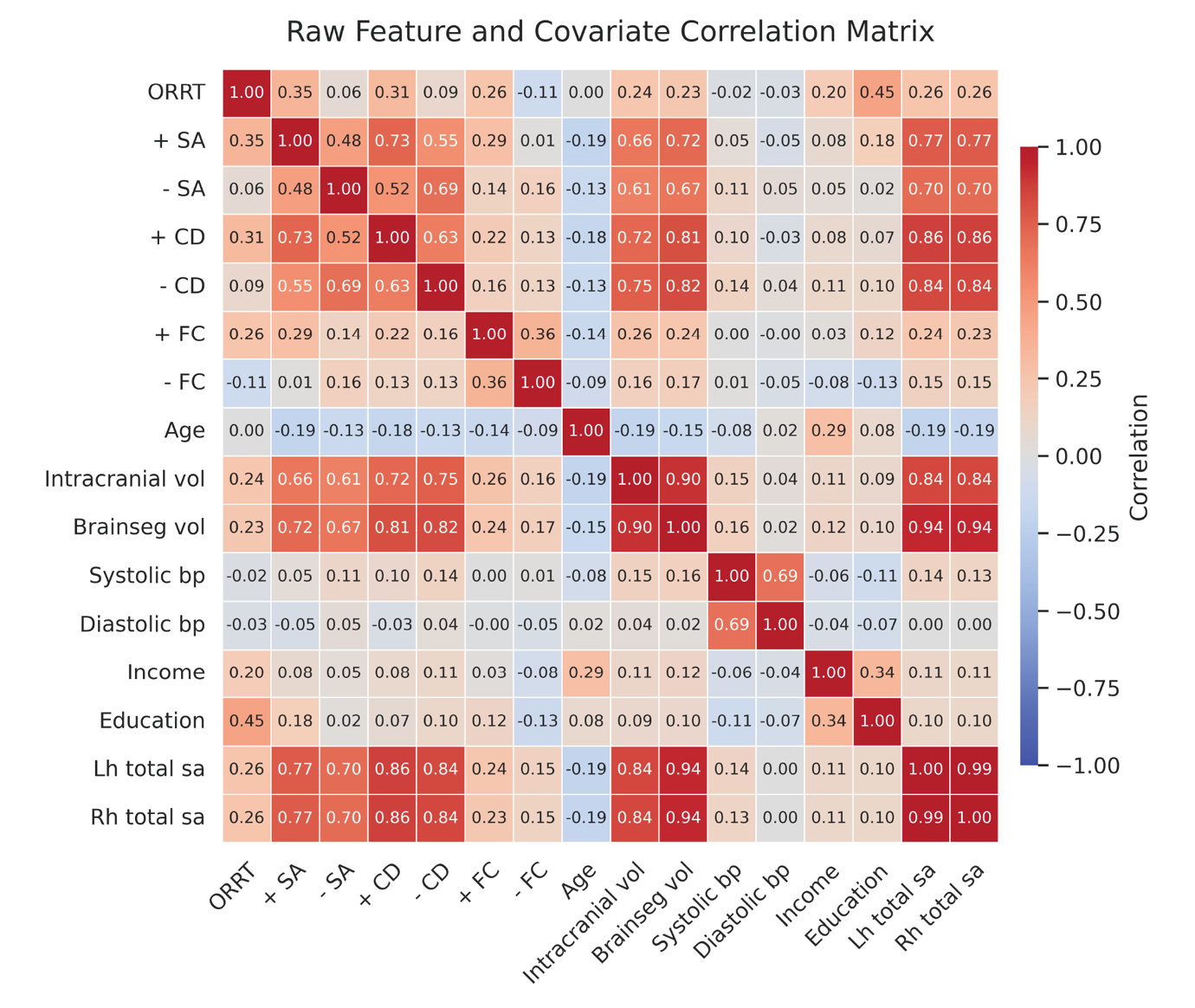

**Fig. S6 | Correlations between behavior, features, and confounds in the discovery cohort.** Raw Pearson correlation matrix from the discovery cohort including word recognition and pronunciation as measured by the Oral Reading Recognition Test (ORRT); the stable positive and negative feature sets derived from the Schaefer 400 parcellation for surface area (SA), cortical distance (CD), and functional connectivity (FC); and confound variables. Confounds: age, intracranial volume, brain segmentation volume, systolic and diastolic blood pressure, income, education, and left and right hemisphere total surface area.

**Supplementary Tables**

|  | **Positive** | | | **Negative** | | | **Joint** | | | **Joint Multimodal** | | | |
| --- | --- | --- | --- | --- | --- | --- | --- | --- | --- | --- | --- | --- | --- |
| **Metric** | **SA** | **CD** | **FC** | **SA** | **CD** | **FC** | **SA** | **CD** | **FC** | **SA + CD** | **SA + FC** | **CD + FC** | **All** |
| **r** | 0.132 | 0.105 | 0.102 | 0.079 | 0.132 | 0.086 | 0.132 | 0.148 | 0.177 | 0.161 | 0.192 | 0.216 | 0.215 |
| **p** | **0.013** | **0.029** | **0.014** | 0.093 | **0.014** | **0.029** | **0.014** | **0.008** | **0.006** | **0.007** | **0.004** | **0.004** | **0.004** |

**Table S1.** Cross-validation model performance predicting word recognition and pronunciation as measured by the Oral Reading Recognition Test performance in the discovery cohort (n = 996). Performance is reported as Pearson's r with corresponding p-values FDR-corrected using the Benjamini–Hochberg procedure. Bold p-values are significant (p < 0.05).
**Abbreviations:** SA, surface area; CD, cortical distance; FC, functional connectivity. Multi-measure labels (e.g., SA+CD) indicate multimodal models combining the joint feature sets of the listed measures; All = SA+CD+FC.

| **Model** | **+ SA** | **+ CD** | **+ FC** | **- SA** | **- CD** | **- FC** | **Joint SA** | **Joint CD** | **Joint FC** | **SA + CD** | **SA + FC** | **CD + FC** | **All** |
| --- | --- | --- | --- | --- | --- | --- | --- | --- | --- | --- | --- | --- | --- |
| **Positive surface area** | — | 0.125 | **0.016** | **0.008** | 1.000 | **0.023** | 1.000 | 1.000 | **0.008** | **0.016** | **0.008** | **0.008** | **0.008** |
| **Positive cortical distance** | 0.125 | — | 1.000 | 1.000 | **0.016** | 1.000 | 0.967 | **0.008** | **0.008** | **0.008** | **0.008** | **0.008** | **0.008** |
| **Positive functional connectivity** | **0.016** | 1.000 | — | 1.000 | 0.679 | 1.000 | 0.351 | **0.008** | **0.008** | **0.008** | **0.008** | **0.008** | **0.008** |
| **Negative surface area** | **0.008** | 1.000 | 1.000 | — | **0.008** | 1.000 | **0.008** | **0.008** | **0.008** | **0.008** | **0.008** | **0.008** | **0.008** |
| **Negative cortical distance** | 1.000 | **0.016** | 0.679 | **0.008** | — | **0.016** | 1.000 | **0.008** | **0.023** | **0.008** | **0.008** | **0.008** | **0.008** |
| **Negative functional connectivity** | **0.023** | 1.000 | 1.000 | 1.000 | **0.016** | — | 0.226 | **0.008** | **0.008** | **0.008** | **0.008** | **0.008** | **0.008** |
| **Joint surface area** | 1.000 | 0.967 | 0.351 | **0.008** | 1.000 | 0.226 | — | 1.000 | **0.023** | **0.008** | **0.008** | **0.008** | **0.008** |
| **Joint cortical distance** | 1.000 | **0.008** | **0.008** | **0.008** | **0.008** | **0.008** | 1.000 | — | 0.702 | **0.031** | **0.016** | **0.008** | **0.008** |
| **Joint functional connectivity** | **0.008** | **0.008** | **0.008** | **0.008** | **0.023** | **0.008** | **0.023** | 0.702 | — | 1.000 | 1.000 | **0.008** | **0.008** |
| **SA + CD** | **0.016** | **0.008** | **0.008** | **0.008** | **0.008** | **0.008** | **0.008** | **0.031** | 1.000 | — | **0.023** | **0.008** | **0.008** |
| **SA + FC** | **0.008** | **0.008** | **0.008** | **0.008** | **0.008** | **0.008** | **0.008** | **0.016** | 1.000 | **0.023** | — | **0.023** | **0.008** |
| **CD + FC** | **0.008** | **0.008** | **0.008** | **0.008** | **0.008** | **0.008** | **0.008** | **0.008** | **0.008** | **0.008** | **0.023** | — | 1.000 |
| **All (SA + CD + FC)** | **0.008** | **0.008** | **0.008** | **0.008** | **0.008** | **0.008** | **0.008** | **0.008** | **0.008** | **0.008** | **0.008** | 1.000 | — |

**Table S2.** Pairwise paired permutation test p-values for cross-validation model performance comparisons in the discovery cohort. P-values were corrected for multiple comparisons using the Benjamini–Hochberg false discovery rate (FDR) procedure across the family of pairwise tests. Bold values indicate statistically significant comparisons after FDR correction (corrected p < 0.05). Because permutation-based p-values are discrete and constrained by the finite number of permutations (10,000), adjusted p-values occur at a limited set of resolution levels, resulting in repeated values across comparisons. Values of corrected p = 1.000 indicate that the adjusted p-value reached the upper bound of the correction procedure. Diagonal cells (model compared to itself) are shaded.
**Abbreviations:** SA, surface area; CD, cortical distance; FC, functional connectivity. Multi-measure labels (e.g., SA+CD) indicate multimodal models combining the joint feature sets of the listed measures; All = SA + CD + FC.

| **Modality** | **Pearson's r** | **p-value** | **95% CI** |
| --- | --- | --- | --- |
| **Unimodal models** | | | |
| Positive SA | 0.188 | **0.001** | [0.085, 0.280] |
| Positive CD | 0.017 | 0.324 | [-0.080, 0.120] |
| Positive FC | 0.115 | **0.019** | [0.003, 0.220] |
| Negative SA | -0.046 | 0.183 | [-0.150, 0.059] |
| Negative CD | 0.132 | **0.009** | [0.041, 0.210] |
| Negative FC | 0.087 | **0.048** | [-0.008, 0.180] |
| Joint SA | 0.135 | **0.009** | [0.036, 0.220] |
| Joint CD | 0.101 | **0.028** | [0.003, 0.190] |
| Joint FC | 0.164 | **0.003** | [0.050, 0.270] |
| **Multimodal models** | | | |
| Positive SA + Negative CD | 0.210 | **0.0001** | [0.110, 0.290] |
| Positive SA + Joint FC | 0.209 | **0.0001** | [0.090, 0.310] |
| Negative CD + Joint FC | 0.207 | **0.0003** | [0.100, 0.309] |
| All | 0.230 | **0.0002** | [0.120, 0.330] |

**Table S3.** Model generalization for predicting word recognition and pronunciation as measured by the Oral Reading Recognition Test performance in the external validation cohort. Results are shown for unimodal models (each based on a single measure) and multimodal models combining the best-performing unimodal feature sets. P-values are FDR-corrected using the two-step Benjamini–Hochberg procedure applied separately to unimodal and multimodal model families. Bold p-values are significant (p < 0.05).
**Abbreviations:** SA, surface area; CD, cortical distance; FC, functional connectivity; CI, confidence interval.

| **Modality** | **Pearson's r** | **p-value** | **95% CI** |
| --- | --- | --- | --- |
| **Unimodal models** | | | |
| Positive SA | 0.150 | **0.003** | [0.052, 0.247] |
| Positive CD | -0.020 | 0.695 | [-0.121, 0.083] |
| Positive FC | 0.126 | **0.013** | [0.022, 0.228] |
| Negative SA | -0.004 | 0.926 | [-0.111, 0.101] |
| Negative CD | 0.105 | **0.035** | [0.003, 0.204] |
| Negative FC | 0.048 | 0.348 | [-0.044, 0.137] |
| Joint SA | 0.124 | **0.013** | [0.033, 0.213] |
| Joint CD | 0.064 | 0.210 | [-0.039, 0.166] |
| Joint FC | 0.146 | **0.004** | [0.044, 0.243] |
| **Multimodal models** | | | |
| Positive SA + Negative CD | 0.167 | **< 0.001** | [0.060, 0.260] |
| Positive SA + Joint FC | 0.180 | **< 0.001** | [0.080, 0.278] |
| Negative CD + Joint FC | 0.178 | **< 0.001** | [0.070, 0.275] |
| All | 0.200 | **< 0.001** | [0.097, 0.298] |

**Table S4.** Model generalization to word–picture association ability (Picture Vocabulary Test) in the validation cohort, using models trained on word recognition and pronunciation in the discovery cohort. Results are shown for unimodal models (each based on a single measure) and multimodal models combining the best-performing unimodal feature sets. P-values are FDR-corrected using the two-step Benjamini–Hochberg procedure applied separately to unimodal and multimodal model families. Bold p-values are significant (p < 0.05).
**Abbreviations:** SA, surface area; CD, cortical distance; FC, functional connectivity; CI, confidence interval.

| **Measure** | **+ SA** | **+ CD** | **+ FC** | **- SA** | **- CD** | **- FC** | **ORRT** |
| --- | --- | --- | --- | --- | --- | --- | --- |
| **Cohen's d** | -0.766 | -0.230 | 0.220 | 0.120 | 0.029 | 0.420 | 0.550 |
| **Wasserstein distance** | 0.470 | 0.120 | 0.260 | 0.080 | 0.043 | 0.410 | 0.520 |

**Table S5.** Cohen's d and Wasserstein distance between the prediction-ready features and reading scores in the HCP-YA discovery cohort and the HCP-A validation cohort. Positive values indicate larger distributions in the discovery cohort.
**Abbreviations:** SA, surface area; CD, cortical distance; FC, functional connectivity; ORRT, Oral Reading Recognition Test.
